# Expression-based selection identifies a microglia-tropic AAV capsid for direct and CSF routes of administration in mice

**DOI:** 10.1101/2024.09.25.614546

**Authors:** Miguel C. Santoscoy, Paula Espinoza, Killian S. Hanlon, Luna Yang, Lisa Nieland, Carrie Ng, Christian E. Badr, Suzanne Hickman, Demitri de la Cruz, Ana Griciuc, Joseph Elkhoury, Rachel E. Bennett, Shiqian Shen, Casey A. Maguire

## Abstract

Microglia are critical innate immune cells of the brain. *In vivo* targeting of microglia using gene-delivery systems is crucial for studying brain physiology and developing gene therapies for neurodegenerative diseases and other brain disorders such as NeuroAIDS. Historically, microglia have been extremely resistant to transduction by viral vectors, including adeno-associated virus (AAV) vectors. Recently, there has been some progress demonstrating the feasibility and potential of using AAV to transduce microglia after direct intraparenchymal vector injection. Data suggests that combining specific AAV capsids with microglia-specific gene expression cassettes to reduce neuron off-targeting will be key. However, no groups have developed AAV capsids for microglia transduction after intracerebroventricular (ICV) injection. The ICV route of administration has advantages such as increased brain biodistribution while avoiding issues related to systemic injection. Here, we performed an *in vivo* selection using an AAV peptide display library that enables recovery of capsids that mediate transgene expression in microglia. Using this approach, we identified a capsid, MC5, which mediated enhanced transduction of microglia after ICV injection compared to AAV9. Furthermore, MC5 enhanced both the efficiency (85%) and specificity (93%) of transduction compared to a recently described evolved AAV9 capsid for microglia targeting after direct injection into the brain parenchyma. Exploration of the use of MC5 in a mouse models of Alzheimer’s disease revealed transduced microglia surrounding and within plaques. Overall, our results demonstrate that the MC5 capsid is a useful gene transfer tool to target microglia *in vivo* by direct and ICV routes of administration.

## Introduction

Microglia, resident myeloid-lineage cells of the central nervous system, are implicated in central nervous system (CNS) pathologies involving neuroinflammation, including Alzheimer’s Disease (AD), AIDS, and brain tumors. The ability to genetically modulate microglia would have a significant impact on the ability to treat these neuroinflammatory diseases. However, they have been recalcitrant to transduction by viral vectors, including adeno-associated virus (AAV) vectors. This may be due to several factors, including the phagocytic nature of these immune cells and the ability to degrade pathogens such as viruses. Another challenge of microglia transduction may relate to the natural neuronal tropism of AAV and the relatively low percentage of microglia (∼10%) compared to other cell types in the brain. The combination of this tropism “sponge” towards neurons and unfavorable stoichiometry may impact whether a given AAV particle is able to interact with microglia. Fortunately, in the past two years, there has been some progress demonstrating the feasibility and potential of using AAV to transduce microglia after direct intraparenchymal vector injection(1, 2). Data suggests that combining specific AAV capsids with microglia-specific gene expression cassettes to reduce neuron off-targeting will be key(1). However, to the best of our knowledge, no groups have developed AAV capsids for microglia transduction via intracerebroventricular (ICV) injection. The ICV route of administration has advantages such as increased brain biodistribution while avoiding issues related to systemic injection. Here, we developed and validated a variant of the AAV transgene-expression selection system, iTransduce(3), combined with an AAV9 peptide display library for microglia transduction in mice after ICV and direct intraparenchymal brain injection.

## Results

### *In vivo* AAV peptide library selection identifies enriched capsids that target microglia after ICV injection

We performed an *in vivo* selection using our published iTransduce AAV9 peptide display system (3), to isolate AAV capsids that can transduce microglia after ICV injection. First, we replaced the broadly active Chicken Beta Actin (*CBA)* promoter with a *CD68* (myeloid cell-selective promoter) driving Cre (**Fig. 1a**). This allows expression of Cre in myeloid-derived cells and limits expression in other cells readily transduced by AAV (e.g. neurons). We then performed two rounds of selection. For the first round of selection, we ICV injected the AAV9 peptide display library into two *CX_3_CR-1^GFP^* mice and 5 weeks later, harvested mouse brain and flow sorted GFP-positive microglia. We performed PCR on DNA isolated from the sorted microglia to amplify the 21 bp-containing insert region of the cap gene. Next, we cloned the recovered inserts into the library backbone and produced the library for round two of selection. For round two of selection we performed a selection to identify capsids capable of transducing microglia. We crossed *Ai9-loxP-STOP-tdTomato* mice with *CX_3_CR-1^GFP^* mice to breed *CX_3_CR1^GFP^* x *Ai9* mice (**Fig. 1b**). These mice have GFP+ microglia and when Cre is expressed by an AAV capsid packaging the CD68-Cre expression cassette, the cells will fluoresce tdTomato+ which can be flow-sorted. *CX_3_CR1^GFP^* x *Ai9* mice were injected ICV with the round 2 library and two weeks later, brain was dissociated and GFP^+^/tdTomato^+^ microglia were flow sorted. We also collected the GFP^+^/tdTomato^-^ fraction to assess the peptide profile in this population. Capsid DNA was rescued by PCR amplification and next generation sequencing (NGS) performed to analyze the diversity of 7mer peptide inserts. NGS in round 1 revealed a very large enrichment of peptides over the original unselected library which for some variants was over 1,000-fold. There was an up-to 3.74-fold enrichment of specific peptides from round 1 to round 2. We next chose candidate peptides from the NGS data, based on the highest frequency of variants recovered from GFP+ tdTomato+ cells and GFP+ tdTomato-cells. We chose one candidate peptide for further testing: IRENAQP (name=MC5). MC stands for “microglia capsid.” This peptide was in the top 5 peptides in the following categories: 1) enrichment between rounds 1 and 2 (2.3-fold), 2) highest percentage in tdT+ cells (5%), and 3) highest tdT/GFP ratio (0.05). A peptide database search revealed that MC5 (IRENAQP) shared high homology, 86%, with a motif within mouse syndecan 4 (SDC4), IPENAQP (**Figure 2**). The residues in the same region of human SDC4 are 43% (3/7 residues) homologous to the MC5 peptide and 57% homologous to murine SDC4.

**Figure 1.**
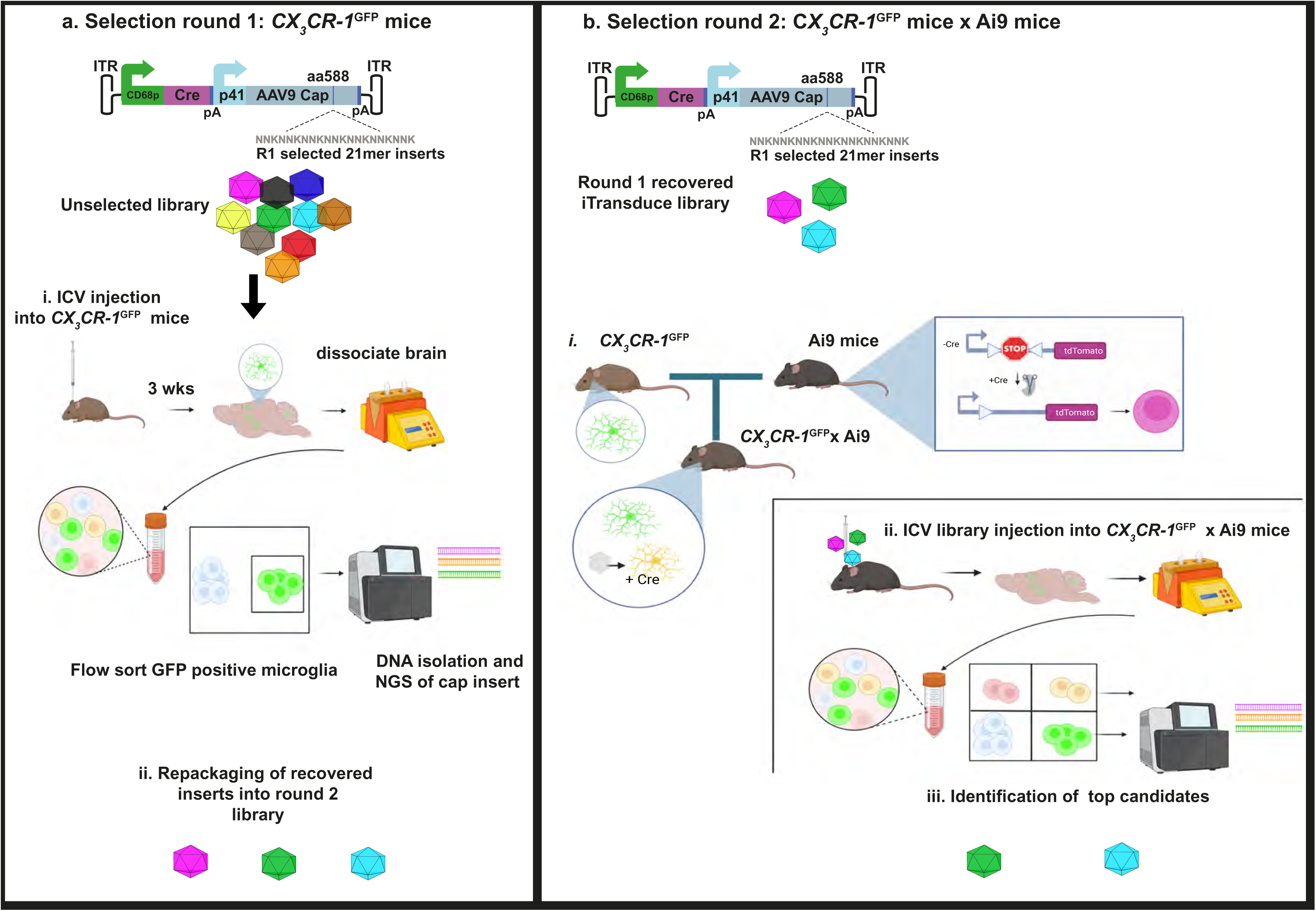
Overview of microglia-transducing AAV capsid selection method. **a.** Round 1 selection. The iTransduce library selection cassette contains a CD68 promoter driving Cre and an AAV p41 promoter driving AAV9 capsid with randomized 21-mer inserts (7-mer peptides) after amino acid 588. This ITR flanked cassette is packaged into the peptide display library. i. *CX_3_CR1^GFP^*mice which express GFP in microglia are injected ICV with the unselected library and 3 weeks later brain is dissociated and GFP^+^ microglia are flow sorted. The capsid gene region flanking the peptide inserts is PCR amplified, analyzed by NGS. ii. The amplified inserts are then repackaging into a library from round 2. **b.** Round 2 selection. In the second round we use the CD68-driven Cre to select capsids that transduce microglia. i. *CX_3_CR1^GFP^* mice are crossed with Ai9 mice. Microglia are GFP^+^ and capsids that express Cre, induce tdTomato expression (microglia express both GFP and tdTomato). ii. Mice are injected ICV with the condensed library from round 1 and brains dissociated 3 weeks later. Transduced microglia (double positive) and GFP^+^ microglia are flow sorted and NGS performed on both populations. iii. Candidate peptides are chosen from these data.

**Figure 2.**
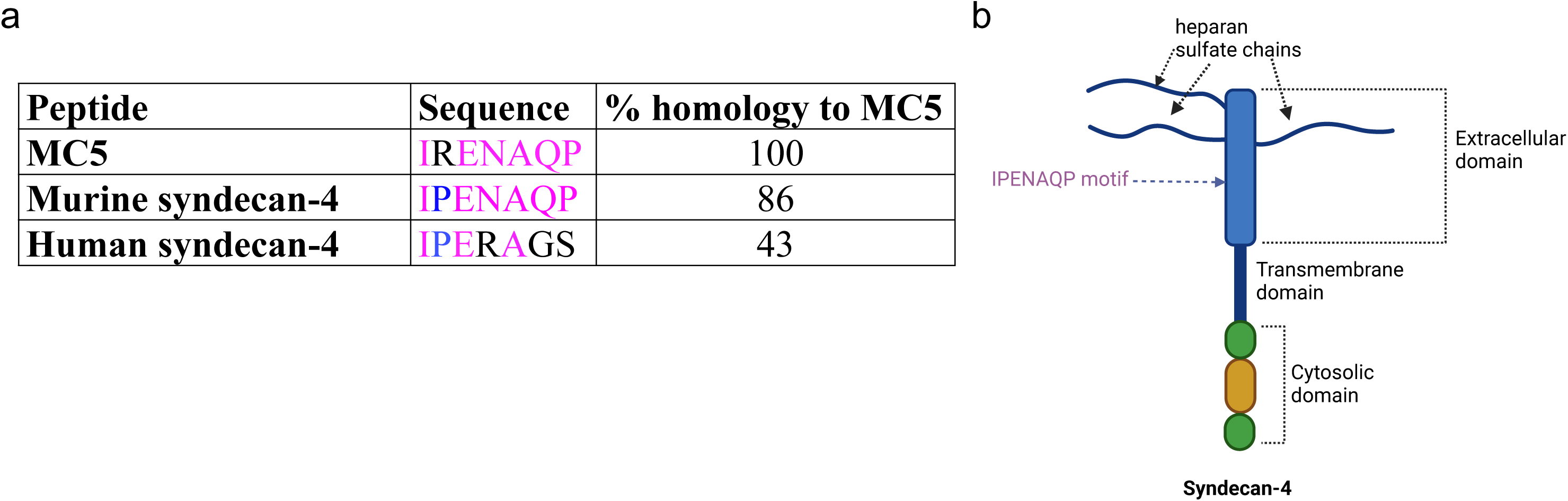
The MC5 capsid’s 7-mer peptide is a putative syndecan-4 motif. **a.** Alignment of the MC5 amino acids (aa) with murine and human syndecan-4. Conserved aa’s are shown in magenta, identity between murine and human only in blue, and non-conserved residues in black. **b.** Schematic of murine syndecan-4 depicting key domains as well as the IPENAQP motif with high identity to the MC5 peptide.

### MC5 mediates higher transduction efficiency than AAV9 after ICV injection in mice

Our next objective was to compare the ability of MC5 vs AAV9 (the parental capsid) to transduce microglia after ICV injection in adult mice. The nucleotide sequences encoding IRENAQP were individually cloned into an AAV9 rep/cap plasmid after amino acid 588 of VP1. For the transgene expression cassette, we used the pAAV-Iba1-GFP-miR9T-miR129-2-3pT construct from Okada et al, which allows for microglia transduction(1)(**Fig. 3a**). Adult female C57BL/6 mice (n=5/capsid) were injected bilaterally with 1.6×10^10^ vg/ventricle of each capsid. One week post injection mice were euthanized and brains harvested for cryosectioning and immunofluorescence staining for GFP and for the microglia marker, Iba1. For both groups, we observed intense immunostaining for GFP which co-localized with Iba1 immediately around the ventricles (**Fig. 3b**). GFP+/Iba1+ cells were also observed in the corpus callosum and cortex near the ventricles (**Fig. 3b**). Next, we performed quantitation of the percentages of Iba1+ microglia transduction by each capsid in the cortex and area surrounding the ventricle in both groups. For each animal, we analyzed five sections adjacent to the ventricle. AAV9 transduced an average of 5.2% (range 2.5-9.9%) of Iba1+ microglia while MC5 transduced an average of 20.7% (range 9.6-63.55%) of Iba1^+^ microglia, a 3.98-fold increase (p<0.015, **Fig 3d**).

**Figure 3.**
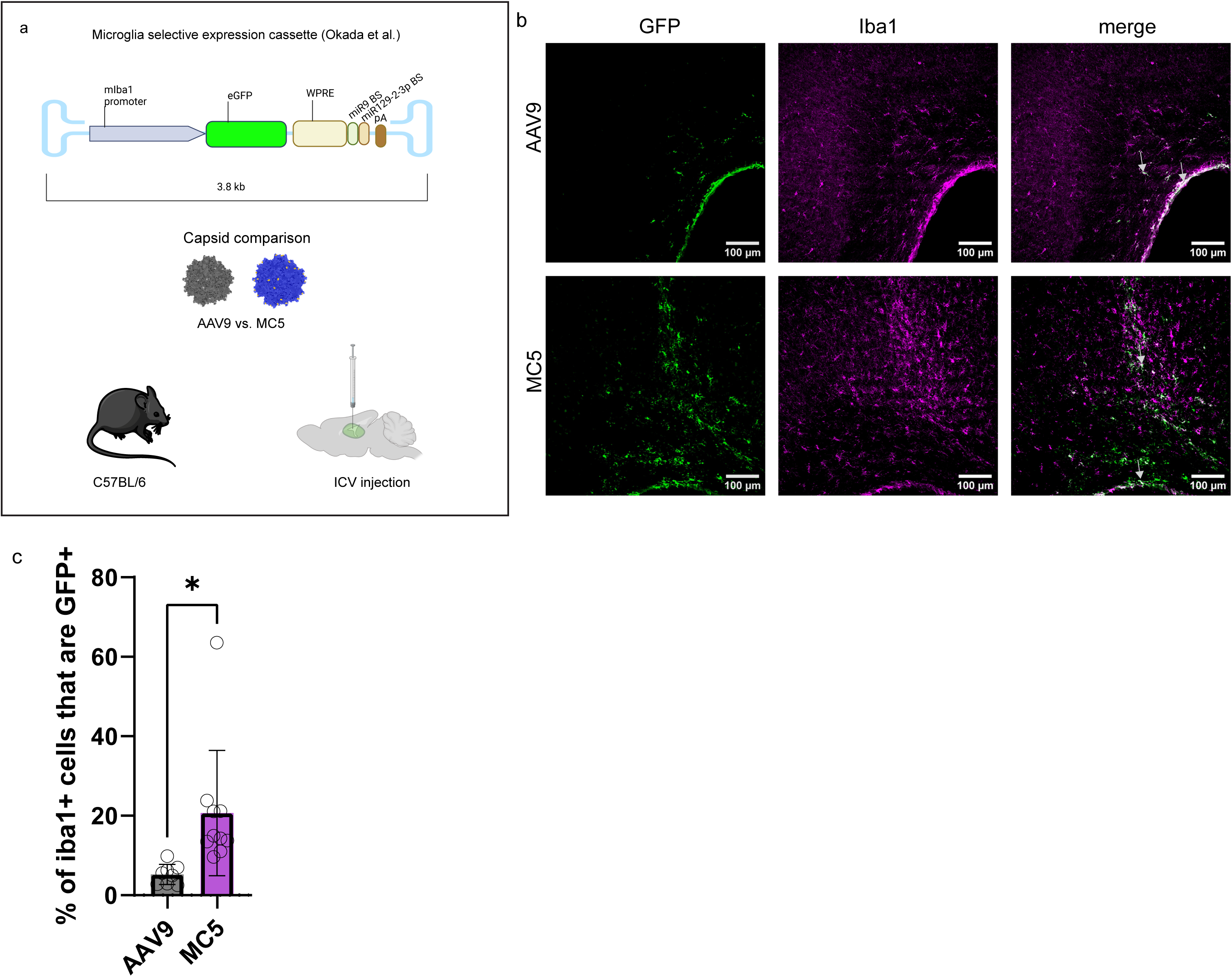
The MC5 capsid is more efficient than the parental AAV9 capsid at transduction of microglia after ICV injection in adult mice. **a.** Schematic of the experiment. The microglia selective transgene expression cassette from Okada et al. was utilized and packaged into AAV9 or MC5. Adult C57BL/6 mice (n=4-5 per group) were injected ICV with either vector. **b.** Confocal imaging surrounding the lateral ventricle to detect vector transduction of cells (GFP, green) and microglia (Iba1, magenta). Arrows point to representative transduced microglia in brain parenchyma and lining the ventricles. **c.** Transduction efficiency of microglia for each capsid. Individual brain sections containing the lateral ventricle (two per mouse) are shown as individual data points. Error bars represent standard deviation of the mean. *p=0.015.

### MC5 mediates efficient transduction of microglia after direct injection into brain parenchyma

Recently an AAV9-based capsid displaying a unique 7-mer peptide called MG1.2 was demonstrated to transduce microglia after direct intraparenchymal injection in mice(2). Here we assessed the specificity and transduction efficiency of MC5 compared to AAV9 and MG1.2 all packaging the AAV-Iba1-GFP-miR9T-miR129-2-3pT genome after intra-hippocampus injection in adult C57Bl/6 mice. Based on the results of Okada et al. which demonstrated that specificity of transduction of microglia with AAV9-Iba1-GFP-miR9T-miR129-2-3pT genome was dose dependent, we tested three doses, (2.1×10^9^ vg, 1.0×10^9^ vg, 0.52×10^9^ vg) injected in a 1.2µl volume in the hippocampus (n=3 mice/dose/capsid). Mice were killed 22 days post injection and intrinsic GFP was imaged along with Iba1 immunostaining by fluorescence microscopy. We quantitated the percentages of GFP+ microglia as well as the percentages of non-microglia such as neurons transduced by each capsid for the 2.1×10^9^ vg, 1.0×10^9^ vg doses. We also compared MC5 transduction efficiency and specificity at all three doses. At the highest dose with all capsids, MC5 had approximately 3-fold more GFP positive microglia (41.3%) than AAV9 (13.3%) or MG1.2 (12.6%) (**Fig. 4a,b**.). MC5 also enabled greater selectivity (∼3-4 fold) over non-microglial cells with 58.2% of transduced cells being microglia vs only 19.6% and 14.3% for AAV9 and MG1.2, respectively (**Fig. 4a, c**). Transduced neurons in the CA1 region of the hippocampus were detected for all capsids although it was less pronounced for MC5 (**Fig. 4a**). The increased specificity was reflected in MC5 transducing the lowest percentage of Iba1^-^GFP^+^ out of total GFP^+^ cells as compared to AAV9 and MG1.2 capsids (**Fig. 4d**). In contrast, at the mid dose, 1.0×10^9^ vg, the highest number of GFP positive cells were observed co-labeling with Iba1^+^ microglia for all capsids. Transduced Iba1+ microglia were observed for AAV9, however intense labeling of neurons in the CA1 region was also observed (**Fig. 5a**). Furthermore, less neuronal transduction was observed for MC5 and MG1.2, suggesting higher specificity of these capsids compared to AAV9 (**Fig. 5a**). High magnification imaging of the transduced microglia showed typical microglia morphology with fine processes clearly visible (**Fig. 5b**). The quantitation of the percentage of transduced microglia revealed that MC5 had the highest (85%), followed by MG1.2 (47%) and AAV9 (32%) (**Fig. 5c**). We measured the specificity of microglia transduction for each capsid by measuring the percentage of GFP^+^ microglia over all GFP positive cells. Remarkably, 93% of GFP^+^ cells transduced by MC5 were microglia which was significantly higher as compared to MG1.2 (83%) and AAV9 (43%) (**Fig. 5d**). We compared the percentages of GFP cells in either the Iba1^+^ or Iba1^-^ (e.g. neurons) cell populations. MC5 had the highest on-target specificity followed by MG1.2 and then AAV9 (**Fig. 5e**). The comparison of MC5 to itself at the three doses revealed a clear benefit of the mid dose for both the highest % of transduced microglia as well as selectivity over non-microglia cells (**Fig. 6**).

**Figure 4.**
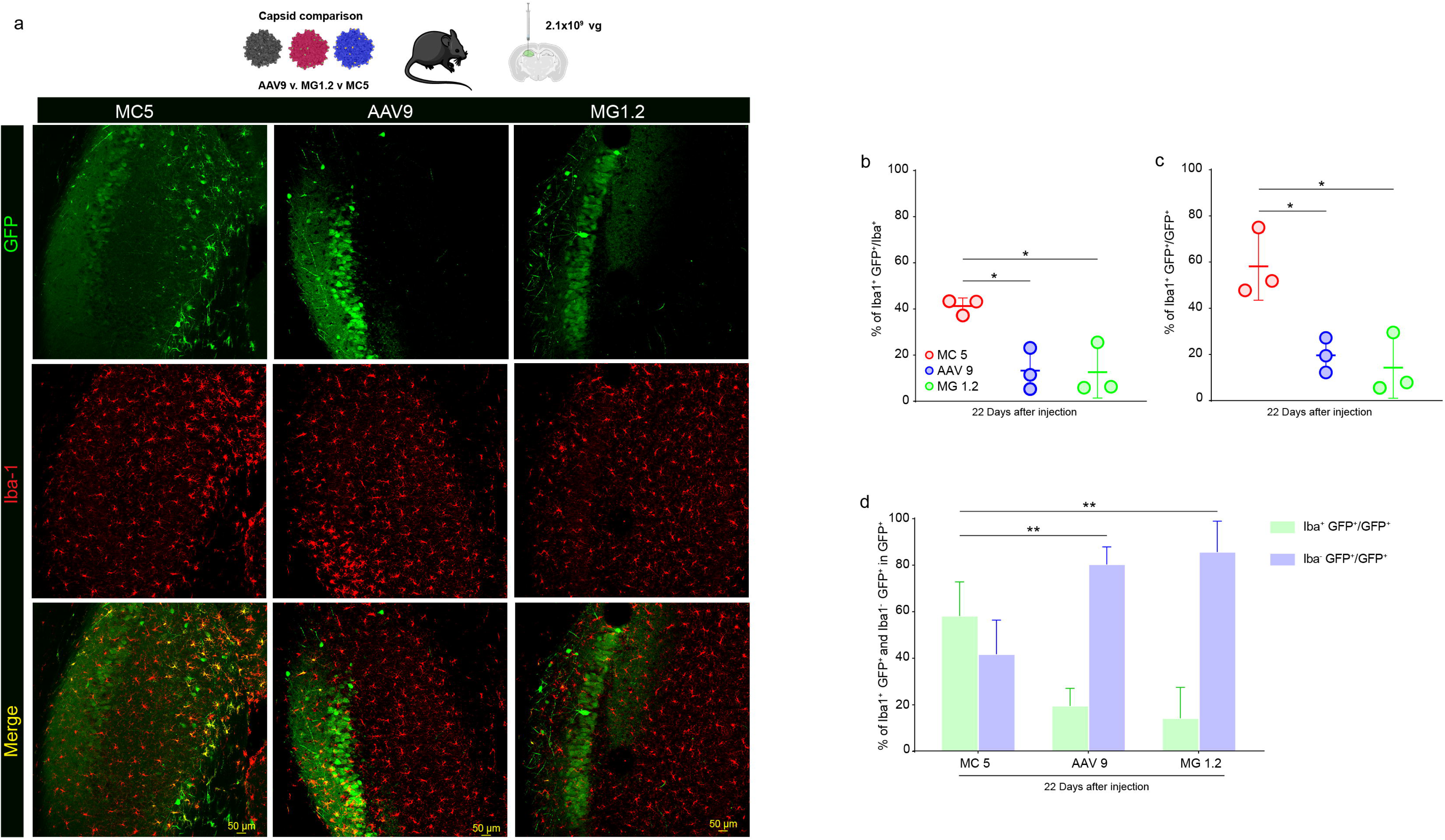
Transduction of microglia by MC5, AAV9, and MG1.2 in the hippocampus after direct injection of the highest dose tested (2.1×10^9^ vg). **a.** Immunofluorescence detection of AAV capsid transduction (GFP) and microglia (Iba1) in mice injected into the hippocampus (n=3 mice/group). GFP is shown in green and Iba1 in red. Colocalization is visualized as yellow in the merge image. Scale bar= 50 µm. **b.** Transduction efficiency of microglia in the hippocampus by each capsid. MC5 vs. AAV9 *,p=0.0216; MC5 vs. MG1.2 *,p= 0.0192. **c.** Transduction specificity for microglia of each capsid. MC5 vs. AAV9, *, p=0.0248; MC5 vs. MG1.2, *, p= 0.0137. **d.** Transduction specificity of microglia (Iba1+) and other cells (Iba1-) cells for each capsid in the hippocampus. MC5 vs. AAV9,**, p=0.0067; MC5 vs. MG1.2, **, p=0.0026.

**Figure 5.**
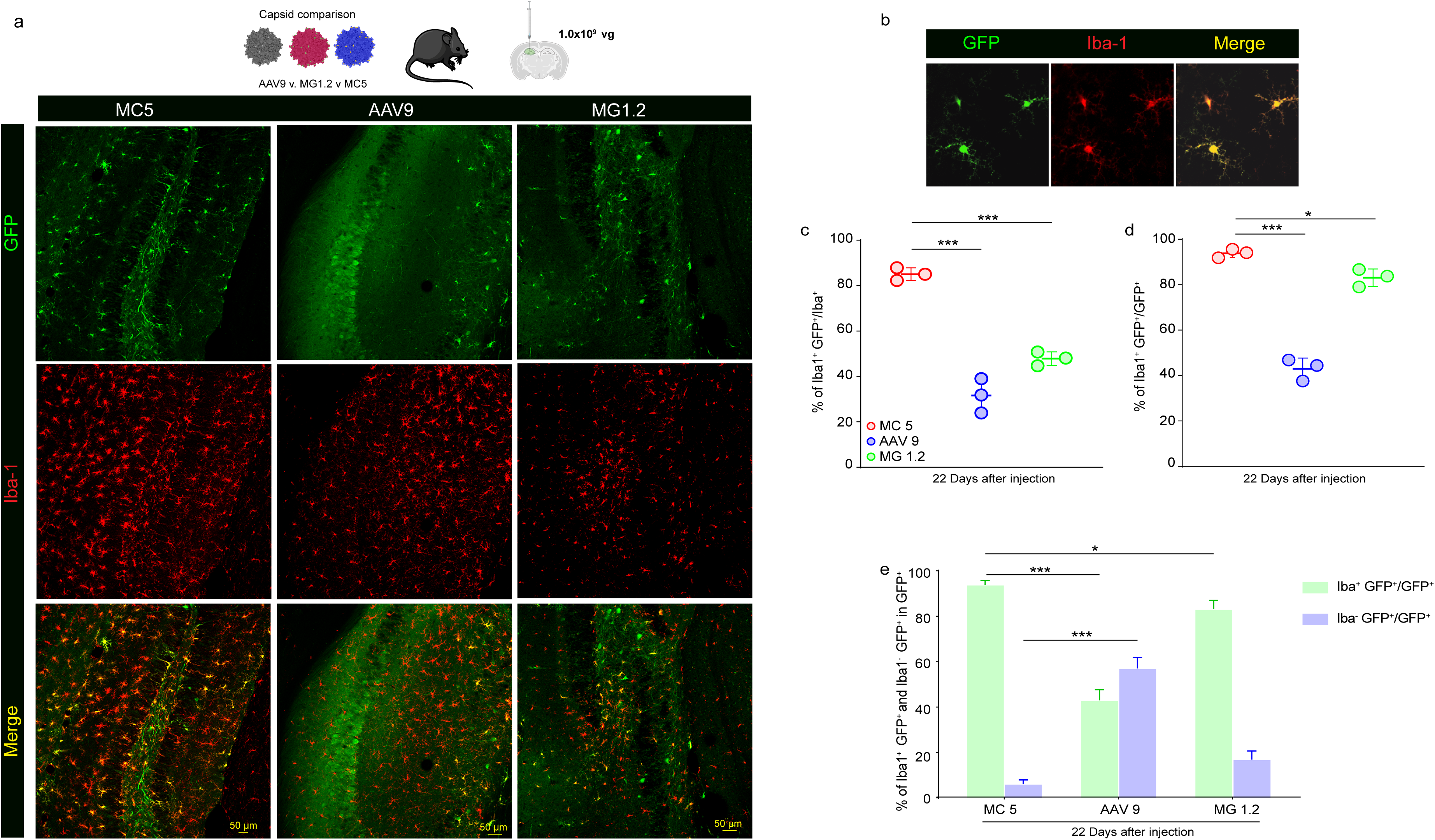
MC5 mediates enhanced transduction efficiency and specificity towards microglia after intracranial injection in hippocampus (1.0×10^9^ vg). **a.** Immunofluorescence detection of AAV capsid transduction (GFP) and microglia (Iba1) in mice injected into the hippocampus (n=3 mice/group). GFP is shown in green and Iba1 in red. Colocalization is visualized as yellow in the merge image. Scale bar= 50 µm. **b.** High magnification of MC5-transduced microglia. **c.** Transduction efficiency of microglia in the hippocampus by each capsid. ***,p=0.001. **d**. Transduction specificity for microglia of each capsid. *,p=0.0363; ***, p=0.001. **e.** Transduction specificity of microglia (Iba1+) and other cells (Iba1-) cells for each capsid in the hippocampus. *,p=0.012; ***, p=0.001.

**Figure 6.**
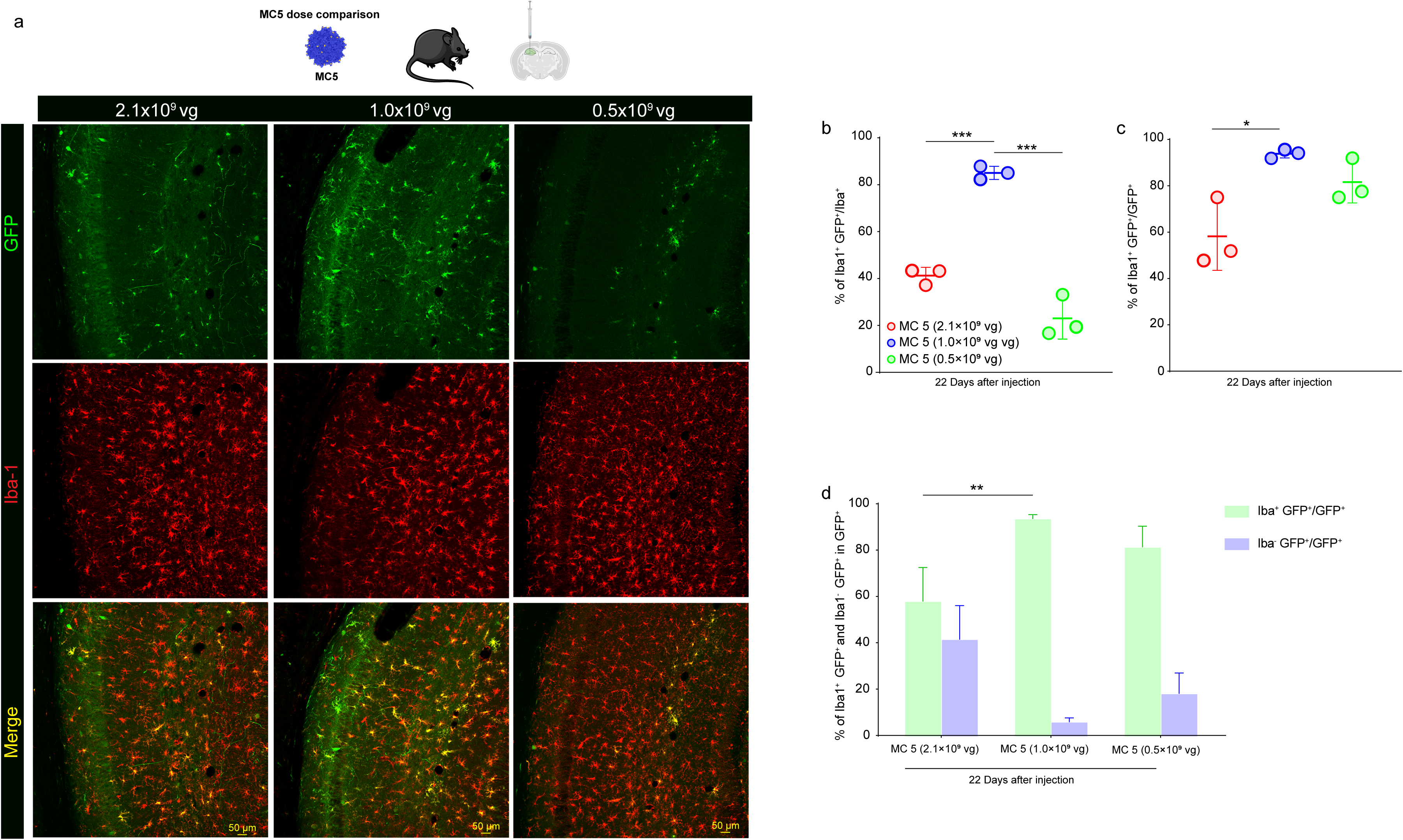
Transduction of microglia by MC5 in the hippocampus after direct injection at three tested doses. **a.** Immunofluorescence detection of AAV capsid transduction (GFP) and microglia (Iba1) in mice injected into the hippocampus (n=3 mice/group). GFP is shown in green and Iba1 in red. Colocalization is visualized as yellow in the merge image. Scale bar= 50 µm. **b**. Transduction efficiency of microglia in the hippocampus by MC5. ***, p=0.001. **c.** Transduction specificity for microglia of MC5. *, p= 0.0142. **d.** Transduction specificity of microglia (Iba1+) and other cells (Iba1-) cells for MC5 in the hippocampus. **, p=0.0028.

### MC5 transduces microglia in APP/PS1 mice with amyloid **β** plaques

To explore whether MC5 could transduce microglia in a mouse model of AD, APP/PS1 mice were injected into the cortex with 8.3×10^8^ vg of MC5-AAV-Iba1-GFP-miR9T-miR129-2-3pT. One week later, mice were sacrificed, and brains sectioned, labeled for amyloid beta (Aβ), and imaged by confocal microscopy. We observed GFP^+^ microglia in the cortex, some of which were surrounding or within Aβ plaques (**Fig. 7**).

**Figure 7.**
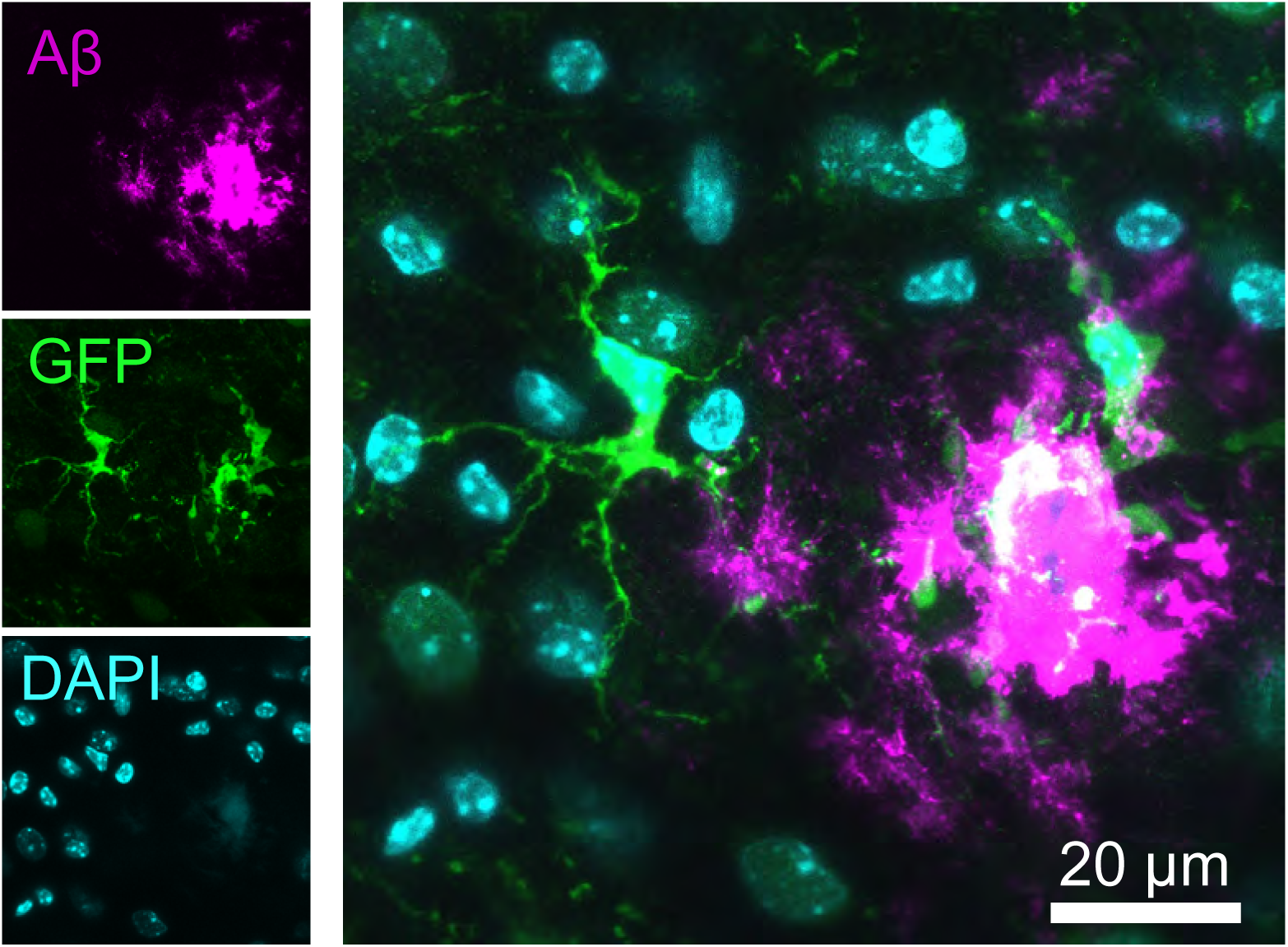
MC5 transduces microglia in *APP/PS1* mice with amyloid β plaques. At 7 days post-injection, GFP-positive plaque-associated microglia were observed throughout cortex in 8-month-old APP/PS1 mice. Image is an 8-micron thick z-projection image. Scale bar= 20 µm.

## Discussion

In this study we set out to select for AAV capsids from a peptide display library with enhanced transduction of microglia. We chose to target microglia via injection into cerebral spinal fluid (CSF) over systemic injection. Direct CSF injection requires far lower dosing compared to the intravenous route and also avoids systemic exposure of vector to the peripheral immune system which has led to severe adverse events (SAEs) in some clinical trials using high-dose AAV vectors (4–7). The concentrations of anti-AAV antibodies are also generally lower in the CSF than the blood, which increases the number of patients eligible for dosing(8). Compared to direct intraparenchymal injection, CSF injection leads to greater vector dispersion, although deeper brain structures such as the striatum are not transduced as efficiently(9). We compared MC5 with AAV9 for transduction of microglia after lateral ventricle injection. MC5 improved the efficiency by ∼4-fold over AAV9 and most transduced microglia were observed lining the ventricles and in the proximal areas of the corpus callosum and cortex. This was performed with one dose and at 7 days post injection. In the future testing different doses and extending the in-life period out to several weeks may improve the detection of more transduced microglia throughout the brain.

Interestingly, in addition to its enhanced transduction via ICV injection, MC5 was efficient at transduction of microglia after direct intraparenchymal injection in the hippocampus. In a recent study, Lin et al. used a directed evolution approach to select AAV capsids that could transduce microglia in mice after direct intracranial injection(2). While very promising, the capsids MG1.1 and MG1.2 were shown to transduce microglia in transgenic mice (e.g. *CX3cr1^CreER^*) that expressed Cre only in microglia and in which the AAV transgene was Cre-inducible (AAV-SFFV promoter-DIO-mScarlet). When MG capsids packaging the Cre inducible reporter were co-injected with AAV packaging a Cre cassette under control of a strong promoter, neurons and astrocytes were transduced by this capsid (and not microglia). Thus, while MG capsids seem to be valuable tools to study microglia biology in transgenic mice, their use as a therapeutic delivery vehicle that can selectively transduce microglia was currently untested. A study by Okada *et al.* demonstrated that AAV9 can transduce microglia after direct intracranial injection in mice if the transgene expression cassette is designed with a microglia selective promoter (*Iba1*) combined with miRNA seed sequences (pAAV-Iba1-GFP-miR9T-miR129-2-3pT) that allow degradation of vector expressed transgene mRNA in non-target cells (e.g. neurons)(1). They found that microglia-selective transduction was dose dependent and increasing the dose changed the profile to primarily neuronal transduction. In our current study, we performed a head-to-head comparison of AAV9, MC5, and MG1.2 capsids all packaging the Okada et al. pAAV-Iba1-GFP-miR9T-miR129-2-3pT genome and injected them in parallel directly into the murine hippocampus. We confirmed the dose-dependent results of Okada et al. that doses above a certain threshold yield significant neuronal transduction and lowering the dose was required for selective microglia transduction for all capsids (**Figs. 4-6**). MC5 was more selective and had higher transduction efficiency of microglia compared to both AAV9 and MG1.2, and at the optimal dosed reached over 80% transduction efficiency and 90% specificity for microglia (**Figs. 5, 6**). These data provide evidence that both physical targeting and transgene expression cassette design (i.e. transcriptional targeting) are important in obtaining the most efficient and selective AAV capsids for *in vivo* microglia transduction.

Based on Okada et al. data, it was not surprising that at the highest dose tested there was more neuronal transduction by MC5 compared to the lower doses tested (**Fig. 4c,d)**. However, what was intriguing was that microglia transduction efficiency was doubled when decreasing the dose of MC5, which would initially seem counterintuitive (**Fig. 6b**). This may indicate that at higher doses, AAV capsids may activate microglia leading to either transcriptional shutdown of transgene expression or degradation of capsids and/or vector genomes. In fact, in pilot studies with high titer, undiluted stocks of MC5, we observed transduced microglia with an ameboid shape, which is an indication of activation (data not shown). There is evidence that suggests that AAV genomes stimulate a TLR9-dependent activation of cytokine release in innate immune cells such as plasmacytoid dendritic cells which is driven by CpG motifs in the AAV vector(10, 11). This can even occur in the brain as it has been reported that intracranially injected AAV can lead to reduce dendritic complexity in transduced neurons and this can be rescued by blocking TLR9 activation with the antagonist oligonucleotide (ODN) 2088(12). As innate immune cells themselves, microglia express TLR9 and activation of this pathway can mediate pro-inflammatory activation (13, 14). Thus, it is quite plausible that at certain dose thresholds, the AAV genome may stimulate TLR9 activation in microglia leading to a variety of effects which may impact AAV mediated transgene expression. For example, inflammatory cytokine release has been shown to reduce transgene expression by AAV vectors(15). In the future, immunosuppressive strategies co-administered with AAV should be tested which may improve transduction efficiency at higher doses, allowing more microglia transduced in larger brain regions.

Syndecans are transmembrane heparan sulfate proteoglycans that interact with a variety of ligands, including integrins, EGFR, and HER2(16). Interestingly, the region of syndecan-4 (SDC4) that the MC5 likely mimics is within the extracellular domain sometimes called the “cell binding domain” as it allows attachment of several cell types (17, 18) (**Fig. 2**). This region, including the NXIPEX motif (part of the region that MC5 has homology to), has been previously identified as highly conserved across mammals(19). Interestingly changing Ile^89^ (contained in the IRENAQP motif of MC5) to alanine in a peptide mimetic of SDC4 reduced SDC4 binding activity to EGFR by 10-fold (19). This motif was also important in binding to α3β1 integrins(19). Since the putative SDC4 motif of the MC5 peptide is in the extracellular region of SDC4, it seems likely that it is engaging a ligand on the surface of microglia, perhaps EGFR and/or α3β1 integrins. It will be interesting in future studies to test MC5 binding to these ligands. The MC5 7-mer ligand may be suitable for affinity maturation/mutagenesis with the aim to improve selectivity of the capsid for microglia. Using the humanized version of the MC5 peptide (**Fig. 2**) as well as the affinity maturation process, we may also be able to develop a translational capsid that may function well *in vivo* in non-human primates and human microglia.

In this study we used the published AAV expression construct by Okada et al. which has an Iba1 promoter, and miR9 and miR129-2-3p target sites. As the field develops, more restrictive promoters and enhancers may be used to further limit expression in neurons. Recently, a preprint described the use of miR124 target sites and the use of a truncated human IBA1 promoter to restrict transduction to microglia(20).

As our experiments with MC5 were done in healthy adult mice, we also wanted to test whether the ability of the capsid to transduce microglia was maintained in relevant disease models. We found in a commonly used mouse model of AD, MC5 transduced microglia surrounding or within Aβ plaques (**Fig. 7**). Thus the MC5 capsid may be useful for studying disease biology and gene therapy strategies targeted at microglia in these models.

Overall, our study demonstrates that the MC5 capsid can be used to transduce microglia in mice and should provide the field with a useful gene delivery vector for preclinical research.

## Materials and Methods

### Cells

293T cells were purchased from American Type Culture Collection (ATCC). Cells were cultured in high glucose Dulbecco’s modified Eagle’s medium containing HEPES (Invitrogen, Carlsbad, CA) supplemented with 10% fetal bovine serum (FBS) (Sigma, St. Louis, MO) and 100 U/mL penicillin, 100 μg/mL streptomycin (Invitrogen) in a humidified atmosphere supplemented with 5% CO2 at 37 °C. Cells were checked regularly for mycoplasma infections using the PCR Mycoplasma Detection Kit (G238; ABM, New York, NY).

### AAV library construction and production

The iTransduce library has been previously described (3, 21). We replaced the broadly active *CBA* promoter with a *CD68* (myeloid cell-selective promoter) driving Cre in the iTransduce plasmid pAAV-CBA-Cre-p41-Cap9 to generate the plasmid pAAV-CD68-Cre-p41-Cap9. Briefly, *pUC57-Cap9-XbaI/KpnI/AgeI* served as template to amplify the AAV9 cap DNA and insert random 21-mer sequences using a forward and reverse primer. Primer information: XF-extend (5’GTACTATCTCTCTAGAACTATTAACGGTTC3’) and reverse primer 588iRev 5’ (GTATTCCTTGGTTTTGAACCCAACCGGTCTGCGCCTGTGCXMNNMNNMNNMNNMN NMNNMNNTTGGGCACTCTGGTGGTTTGTG 3’) in which the MNN repeat refers to the the randomized 21-mer nucleotides (purchased from IDT). The 447 bp PCR product was digested with XbaI and AgeI overnight at 37°C and then we gel-purified the product (Qiagen). Similarly, *pAAV-CBA-Cre-p41-Cap9 or pAAV-CD68-Cre-Cap9* was digested with XbaI and AgeI and gel purified. Next, a ligation reaction (1h at room temperature) with T4 DNA ligase (NEB) was performed using a 3:1 cap insert to vector molar ratio. The subsequent ligated plasmid was called *pAAV-CBA-Cre-p41-Cap9-7mer or pAAV-CD68-Cre-p41-Cap9-7mer* and contained a pool of plasmids with random 7-mer peptides inserted in the cap gene between nucleotides encoding 588 and 589 of AAV9 VP3.

We produced the library as previously described(3). Briefly, 293T cells were transfected using PEI MAX^®^ solution (Polysciences, Warrington, PA) with pAAV-CBA-Cre-p41-Cap9-7mer (Round 1 of selection) or pAAV-CD68-Cre-p41-Cap9-7mer (Round 2 of selection), the adenovirus helper plasmid (pAdΔF6, 26 μg per plate), and rep plasmid (pAR9-Cap9-stop/AAP/Rep, 12 μg per plate) to induce production of AAV. AAV was purified from the cell lysate and polyethylene glycol-precipitated media using iodixanol density-gradient ultracentrifugation. Buffer exchange to PBS was done using ZEBA spin columns (7K MWCO; Thermo Fisher Scientific) and further concentration was performed using Amicon Ultra 100kDa MWCO ultrafiltration centrifugal devices (Millipore). Vectors were stored at −80 °C until use. We quantified AAV genomic copies (vg) in AAV preparations using TaqMan qPCR with ITR-sequence specific primers and probes(22, 23).

### *In vivo* library selection

For the first round, we ICV injected the AAV9 peptide display library into two *CX_3_CR-1^GFP^*male mice (3.05×10^8^ vg for a 9-month old mouse and 6.1×10^8^ vg for a 5 month old mouse). Five weeks later, we harvested mouse brains and flow sorted GFP^+^ microglia. To do this, mice were anesthetized with an overdose of ketamine/xylazine and transcardially perfused with phosphate-buffered saline (PBS). Brains were immediately dissociated using the Miltenyi Neural Tissue Dissociation kit (Miltenyi Biotec, Auburn, CA). We slightly modified the original protocol to remove the excess of myelin while maintaining cell viability. Briefly, we placed every brain in one C tube and added the Miltenyi Enzyme P with PBS. For rapid homogenization, the brain was cut into smaller fragments before running the Miltenyi GentleMACS dissociator (Miltenyi Biotec). After three sequential runs of dissociation, we added the previously diluted Miltenyi Enzyme A into Buffer Y. After incubation at 37 °C for 10 min, we added four volumes of 0.5% w/v BSA dissolved in PBS and transferred the brain suspension through a 100 µm cell strainer. Myelin was rapidly removed with Miltenyi Myelin removal beads and EasySep Magnets (Miltenyi Biotec). After the last step of myelin removal using LS columns (Miltenyi Biotec), the cell suspension was immediately sorted for GFP^+^ microglia setting the gates with a freshly processed brain cell suspension of a C57BL/6J mouse. After sorting, the GFP-positive cells were immediately pelleted by centrifugation, and DNA was extracted using the ARCTURUS PicoPure DNA extraction kit (ThermoFisher). After DNA extraction, the Cap9 DNA flanking the 21mer inserts was amplified using the following primers: Cap9_Kpn/Age_For: 5’-AGCTACCGACAACAACGTGT-3’ and Cap9_ Kpn/Age_Rev: 5’-AGAAGGGTGAAAGTTGCCGT-3’ and Phusion High-Fidelity PCR kit (New England Biolabs). The amplicon was gel purified digested with KpnI, and AgeI and the Cap9 KpnI-AgeI fragments (144 bp) were agarose gel purified before ligation in the pUC57-Cap9-XbaI/AgeI/KpnI plasmid (digested with KpnI and AgeI). The ligation product was transformed into electrocompetent DH5alpha bacteria (New England Biolabs) and the entire transformation was grown overnight in LB-ampicillin medium. pUC57-Cap9-XbaI/AgeI/KpnI plasmid was purified by maxi prep (Qiagen). Plasmid was digested by XbaI/AgeI to release the 447 bp cap fragment which was gel purified and ligated with similarly cut pAAV-CD68-Cre-mut/p41-Cap9-7mer for the next round of AAV library production. For round two we performed a selection to identify capsids capable of transducing microglia. *CX_3_CR1^GFP^* x *Ai9* mice were injected ICV with the round 2 library (10^10^ vg) and two weeks later, brain was dissociated and GFP^+^/tdTomato^+^ microglia were flow sorted. We also collect the GFP^+^/tdTomato^-^ fraction to assess the peptide profile in this population. Capsid DNA was rescued by PCR amplification and next generation sequencing (NGS) performed by the Massachusetts General Hospital DNA Core to analyze the diversity of 7mer peptide inserts. For each round of selection vector DNA corresponding to the insert-containing region was amplified by PCR using either Phusion High-Fidelity enzyme or Q5 polymerase (both from New England Biolabs using Forward primer: 5’-AATCCTGGACCTGCTATGGC-3’, and reverse primer: 5’-TGCCAAACCATACCCGGAAG-3’). PCR products were purified using a QIAquick PCR Purification Kit (Qiagen). Unique barcode adapters were annealed to each sample, and samples were sequenced on an Illumina MiSeq (150bp reads) at the Massachusetts General Hospital Center for Computational and Integrative Biology DNA Core. Approximately 50,000-100,000 reads per sample were analyzed. Sequence output files were quality-checked initially using FastQC (http://www.bioinformatics.babraham.ac.uk/projects/fastqc/) and analyzed on a program custom-written in Python. Briefly, sequences were binned based on the presence or absence of insert; insert-containing sequences were then compared to a baseline reference sequence and error-free reads were tabulated based on incidences of each detected unique insert. Inserts were translated and normalized.

### AAV vector production

For transgene expression studies with AAV vectors we used the following AAV expression plasmid:pAAV/mIba1.GFP.WPRE.miR-9.T.miR-129-2-3p.T.SV40pA was a gift from Hirokazu Hirai (Addgene plasmid # 190163; http://n2t.net/addgene:190163; RRID:Addgene_190163)(1). This plasmid was purified by Alta Biotech (Aurora, CO). The plasmid was digested with SmaI restriction enzyme (New England Biolabs, Ipswich, MA) at room temperature for one hour to confirm ITR integrity. Oxford Nanopore complete plasmid sequencing was performed by the MGH DNA Core to confirm plasmid sequence integrity.

We used the following three capsids for these studies: AAV9 which was encoded in the pAR9 rep/cap vector kindly provided by Dr. Miguel Sena-Esteves at the University of Massachusetts Medical School, (Worcester, MA). The MG1.2 capsid is a previously described engineered AAV9-based capsid(2). rAAV2/MG1.2 was a gift from Minmin Luo (Addgene plasmid # 184541; http://n2t.net/addgene:184541; RRID:Addgene_184541). MC5 was generated by digesting pAR9 BsiWI and BaeI which removes a fragment flanking the VP3 amino acid 588 site for peptide sequence insertion. Next, we ordered a 997 bp dsDNA fragment from Integrated DNA Technologies (IDT, Coralville, IA), which contains overlapping Gibson homology arms with the BsiWI/BaeI cut AAV9 as well as the 21-mer nucleotide sequence encoding the peptide of interest in frame after amino acid 588 of VP3. Last, we performed Gibson assembly using the Gibson Assembly® Master Mix (NEB, Ipswich, MA) to ligate the peptide containing insert into the AAV9 *rep/cap* plasmid. After transformation into competent bacteria, we picked single colonies and isolated DNA using minipreps (Qiagen). Complete plasmid sequencing was performed to verify the insert sequence at the MGH DNA core.

AAV production was performed as previously described(24). Briefly, 293T cells were triple transfected using PEI MAX^®^ solution (Polysciences, Warrington, PA) with (1) AAV-rep/cap plasmid (either AAV9, MG1.2, or MC5) (2) an adenovirus helper plasmid, pAdΔF6, and (3) ITR-flanked AAV transgene expression plasmid (pAAV/mIba1.GFP.WPRE.miR-9.T.miR-129-2-3p.T.SV40pA). Cell lysates and polyethylene glycol-precipitated media containing vector were harvested 68-72 h post transfection and purified by ultracentrifugation of an iodixanol density gradient. Iodixanol was removed and buffer exchanged to phosphate buffered saline (PBS) containing 0.001% v/v Pluronic F68 (Gibco™, Grand Island, NY) using 7 kDa molecular weight cutoff Zeba™ desalting columns, (Thermo Scientific). Vector was concentrated using Amicon® Ultra-2 100 kDa MWCO ultrafiltration devices (Millipore Sigma). Vector titers in vg/ml were determined by Taqman qPCR in an ABI Fast 7500 Real-time PCR system (Applied Biosystems) using probes and primers to the ITR sequence and interpolated from a standard curve made with a restriction enzyme linearized AAV plasmid. Vectors were pipetted into single-use aliquots and stored at −80°C until use.

### Mice

All animal experiments were approved by the Massachusetts General Hospital Subcommittee on Research Animal Care following guidelines set forth by the National Institutes of Health Guide for the Care and Use of Laboratory Animals. We used adult age (8-10 week old) C57BL/6J (strain # 000664), B6.129P2(Cg)-*Cx3cr1^tm1Litt^*/J (common name *CX_3_CR-1^GFP^*, strain 005582), and B6.Cg-*Gt(ROSA)26Sor^tm9(CAG-tdTomato)Hze^*/J (common name Ai9, strain 007909), all from The Jackson Laboratory, Bar Harbor, ME. We crossed homozygous Ai9 with homozygous *CX_3_CR-1^GFP^* to yield *Ai9: CX_3_CR-1^GFP^* progeny for the round 2 selection process. We also used *APP/PS1* mice (*B6;C3-Tg(APPswe,PSEN1dE9)85Dbo/Mmjax*; Stock 034829-JAX).

### Intracranial injection of AAV vectors

#### Intracerebroventricular injections into the lateral ventricle

Adult mice were anesthetized using isoflurane and analgesia achieved with buprenorphine (0.15 mg/kg) and local scalp administration of lidocaine (5mg/kg). Once deeply anesthetized, mice were placed into a *Just For Mouse* Stereotaxic Frame with an integrated animal warming base (Stoelting, Wood Dale, IL). Adult mice (n=5/group) were stereotactically injected bilaterally into the left and right lateral ventricles at the dose described in the figure legend of each vector preparation in a volume of 5 µl using the following coordinates from bregma in mm: anterior/posterior, AP −0.4; medial/lateral, ML +/-1.0; dorsal/ventral, DV −1.7. Vectors were infused at a rate of 1.0 µl/min using a Quintessential Stereotaxic Injector pump (Stoelting) to drive a gas-tight Hamilton Syringe (Hamilton, NV) attached to a 10 µl 33-gauge NEUROS model syringe (Hamilton, NV). After injection, the needle was left in place for two minutes to allow the vector solution to disperse and not backflow up the cannula. Buprenorphine (0.15 mg/kg) was injected subcutaneously twice a day for two days after the surgery for analgesia. The in-life portion of the study is indicated in the figure legends.

#### Intra-hippocampus vector injection

AAV vectors (AAV9, MC5, MG1.2) were prepared at different concentrations to deliver three doses of each (0.5×10^9^ vg; 1.0×10^9^ vg, MC 2.1×10^9^ vg) in 1.2 µl PBS. C57BL/6 mice (4-month-old, male, n = 3 mice) were anesthetized with oxygenated isoflurane (3% for induction, 1.5% for maintenance) and mounted on a stereotaxic frame. The scalp was prepared using alcohol swabs (BD, US). After 1% lidocaine infiltration, a midline incision was made using mini scissors. Mini-craniotomy was made at the designated coordinates (AP 2 mm, ML 2 mm). Using a thin glass pipette loaded on Nanoject III (Drummond, US), a total volume of 1.2 µl virus was slowly injected into the hippocampus at the depth of 1.5 mm and 2.0 mm. The needle was left *in situ* for 10 minutes after the injection to minimize backflow of virus during needle retraction. Skin was closed using 4-0 polypropylene suture (Oasis, US). Animals were kept on a warm pad and returned to home cage after full recovery from anesthesia.

#### Intracortical vector injection in APP/PS1 mice

Alternatively, intra-cortical injections in APP/PS1 mice, were similarly performed under isoflurane anesthesia using a 33-gauge Hamilton syringe. Mice received 8.3×10^8^ vg of MC5 in 1 µl which was directly injected into the cortex overlying the hippocampus at a depth of 0.3 mm. Post-operative warming and buprenorphine analgesia was performed as described above.

### Immunofluorescence staining and Microscopy and Image analysis

#### ICV injected mice

Mice were deeply anesthetized with an overdose of ketamine/xylazine and transcardially perfused with phosphate buffered saline (PBS) followed by 4% v/v formaldehyde in 1x PBS. Brains were post-fixed in 4% formaldehyde diluted in PBS for 48 h, followed by 30% (w/v) sucrose for cryopreservation for another 48-72 h after which brains were embedded and frozen in Tissue-Tek ® O.C.T. compound (Sakura Finetek USA, Torrance, CA). Coronal floating sections (40 μm) were cut using a NX50 CryoStar Cryostat (Thermo Scientific). After rinsing off the sucrose in PBS, the brain sections were treated for immunofluorescence or mounted on glass slides for imaging.

For immunofluorescence, the cryosections were permeabilized with 0.5% v/v Triton™ X-100 (Millipore Sigma) in PBS for 2 h and blocked with 5% v/v normal goat serum (NGS) in PBS for 1h. Permeabilization and blocking steps were performed while gentle shaking (30 rpm) at room temperature (RT) in 12-well plates. Brain sections with primary antibodies diluted in 1.5% v/v NGS were incubated at 4°C for 24 h on a platform orbital shaker set at 60 rpm. After three washes with PBS, coronal sections and secondary antibodies diluted in 1.5% v/v NGS were incubated for 1h at RT 60 rpm. Three PBS washes were performed prior mounting of stained sections on glass slides for microscopy. Primary antibodies for staining of AAV transduced cells (GFP) and microglia were chicken anti-GFP (GFP-1020, Aves, Davis, CA) and rabbit anti-Iba1 (019-19741, Fujifilm Wako Chemicals USA), respectively, both at working dilutions of 1:100 in 1.5% v/v normal goat serum (NGS). Secondary antibodies were goat anti-chicken Alexa Fluor 488 for GFP (Thermo Scientific) and goat anti-rabbit Alexa Fluor 647 for Iba1 (Thermo Scientific) both at working dilutions of 1:1000 in 1.5% v/v NGS.

Sections were mounted with Vectashield mounting medium with DAPI (Vector Laboratories, Burlingame, CA) and, imaging was performed with a NIKON CSU-W1 spinning disk confocal microscope.

#### Intra-hippocampus injected mice

On day 22 after virus injection, mice were sacrificed and perfused with ice-cold PBS followed by 4% PFA. Mice brains were extracted and fixed in 4% PFA for additional 2 days at 4°C. Brains were sectioned using a vibratome (Leica, VT1000) at 50mm thickness and the slices covering the hippocampus were collected. The slices were blocked with 5% Bovine Serum Albumin (Boston Bioproducts, USA) and permeabilized with 0.5% Triton™ X-100 (Millipore Sigma) in PBS, followed by primary antibodies rabbit anti-Iba1(1:1000) and secondary antibody goat anti-rabbit Cy3 (1:1000, Jackson ImmunoResearch, USA). The brain slices were subsequently mounted onto slides with DAPI and imaged using a confocal microscope (NIKON AXR, Japan). Interest areas were scanned with a 20x objective, data were analyzed using ImageJ (NIH). Both the injector of the vectors, the imager of the sections, and the analyzer were performed in a group blinded fashion.

#### Intra-cortical injected APP/PS1 mice

At 7 days, brains were collected, fixed in 4% paraformaldehyde for 48 hours and then equilibrated in 30% sucrose in PBS. After 24 hours, 40-micron thick tissue sections were collected on a freezing microtome, labeled for amyloid β (1:500, RRID:AB_2797642) overnight at 4C, then rinsed and coverslipped with Fluoromount G with DAPI (Southern Biotech, cat no. 0100-20). Imaging was performed using an Olympus FV3000 confocal and 63x oil immersion lens.

### Statistics

We used GraphPad Prism 9.0 for PC for statistical analysis. To compare means of two groups, we used an unpaired two tailed t-test; p values <0.05 were accepted as significant. For comparison of transduction of AAV9, MG1.2, and MC5 we used a one-way ANOVA followed by a Šídák’s multiple comparisons test.

## Acknowledgments

For generation of the artwork in the Figures, BioRender software was utilized for some of the objects. We thank the Molecular Imaging Center MGH Campus Navy Yard (CNY) for their input in the method development for collecting images and quantifying data.

## Funding

This work was supported by NIH R01 grant DC017117, NIH NCI 1R35CA232103-01, and a Sanofi iAward sponsored research award (to C.A.M.). This work was also supported by a MassCATS award (to C.A.M. and R.E.B). M.C.S. was partially supported by an American Society of Gene and Cell Therapy Underrepresented Population Fellowship Award in Gene and Cell Therapy.

## Author contributions

C.A.M. conceived of the study, performed experiments, analyzed data, and wrote the manuscript. M.C.S, K.S.H, S.S., L.Y., R.E.B., L.N., S.H., and P.E. performed experiments and analyzed data. D.D.L.C., C.N. performed experiments. A.G., J.E., and C.E.B. analyzed data. All authors assisted in reviewing and editing the manuscript.

## Competing interests

C.A.M. has a financial interest in Sphere Gene Therapeutics, Inc., Chameleon Biosciences, Inc., and Skylark Bio, Inc., companies developing gene therapy platforms. C.A.M.’s interests were reviewed and are managed by MGH and Mass General Brigham in accordance with their conflict-of-interest policies. C.A.M., M.C.S., K.S.H., and P.E. have filed a patent application with claims involving the MC5 capsid.

## Data and materials availability

Data are available upon request. The rep/cap plasmid encoding the MC5 capsid will be available at Addgene upon acceptance of the manuscript in a peer-reviewed journal.

## References

1. Okada Y, Hosoi N, Matsuzaki Y, Fukai Y, Hiraga A, Nakai J, Nitta K, Shinohara Y, Konno A, Hirai H. Development of microglia-targeting adeno-associated viral vectors as tools to study microglial behavior in vivo. Commun Biol. 2022;5(1):1224.

2. Lin R, Zhou Y, Yan T, Wang R, Li H, Wu Z, Zhang X, Zhou X, Zhao F, Zhang L, et al. Directed evolution of adeno-associated virus for efficient gene delivery to microglia. Nat Methods. 2022;19(8):976–85.

3. Hanlon KS, Meltzer JC, Buzhdygan T, Cheng MJ, Sena-Esteves M, Bennett RE, Sullivan TP, Razmpour R, Gong Y, Ng C, et al. Selection of an Efficient AAV Vector for Robust CNS Transgene Expression. Mol Ther Methods Clin Dev. 2019;15:320–32.

4. Lek A, Wong B, Keeler A, Blackwood M, Ma K, Huang S, Sylvia K, Batista AR, Artinian R, Kokoski D, et al. Unexpected Death of a Duchenne Muscular Dystrophy Patient in an N-of-1 Trial of rAAV9-delivered CRISPR-transactivator. medRxiv. 2023:2023.05.16.23289881.

5. Wilson JM, Flotte TR. Moving Forward After Two Deaths in a Gene Therapy Trial of Myotubular Myopathy. Hum Gene Ther. 2020;31(13-14):695–6.

6. Philippidis A. Fourth Boy Dies in Clinical Trial of Astellas’ AT132. Hum Gene Ther. 2021;32(19-20):1008–10.

7. Salabarria SM, Corti M, Coleman KE, Wichman MB, Berthy JA, D’Souza P, Tifft CJ, Herzog RW, Elder ME, Shoemaker LR, et al. Thrombotic microangiopathy following systemic AAV administration is dependent on anti-capsid antibodies. J Clin Invest. 2024;134(1).

8. Gray SJ, Nagabhushan Kalburgi S, McCown TJ, Jude Samulski R. Global CNS gene delivery and evasion of anti-AAV-neutralizing antibodies by intrathecal AAV administration in non-human primates. Gene Ther. 2013;20(4):450–9.

9. Nakamura S, Osaka H, Muramatsu SI, Takino N, Ito M, Jimbo EF, Watanabe C, Hishikawa S, Nakajima T, Yamagata T. Intra-cisterna magna delivery of an AAV vector with the GLUT1 promoter in a pig recapitulates the physiological expression of SLC2A1. Gene Ther. 2021;28(6):329–38.

10. Rogers GL, Shirley JL, Zolotukhin I, Kumar SRP, Sherman A, Perrin GQ, Hoffman BE, Srivastava A, Basner-Tschakarjan E, Wallet MA, et al. Plasmacytoid and conventional dendritic cells cooperate in crosspriming AAV capsid-specific CD8(+) T cells. Blood. 2017;129(24):3184–95.

11. Alakhras NS, Moreland CA, Wong LC, Raut P, Kamalakaran S, Wen Y, Siegel RW, Malherbe LP. Essential role of pre-existing humoral immunity in TLR9-mediated type I IFN response to recombinant AAV vectors in human whole blood. Front Immunol. 2024;15:1354055.

12. Suriano CM, Kumar N, Verpeut JL, Ma J, Jung C, Dunn CE, Carvajal BV, Nguyen AV, Boulanger LM. An innate immune response to adeno-associated virus genomes decreases cortical dendritic complexity and disrupts synaptic transmission. Mol Ther. 2024;32(6):1721–38.

13. Benbenishty A, Gadrich M, Cottarelli A, Lubart A, Kain D, Amer M, Shaashua L, Glasner A, Erez N, Agalliu D, et al. Prophylactic TLR9 stimulation reduces brain metastasis through microglia activation. PLoS Biol. 2019;17(3):e2006859.

14. Maatouk L, Compagnion AC, Sauvage MC, Bemelmans AP, Leclere-Turbant S, Cirotteau V, Tohme M, Beke A, Trichet M, Bazin V, et al. TLR9 activation via microglial glucocorticoid receptors contributes to degeneration of midbrain dopamine neurons. Nat Commun. 2018;9(1):2450.

15. Breous E, Somanathan S, Bell P, Wilson JM. Inflammation promotes the loss of adeno-associated virus-mediated transgene expression in mouse liver. Gastroenterology. 2011;141(1):348–57, 57 e1-3.

16. Wang H, Jin H, Beauvais DM, Rapraeger AC. Cytoplasmic domain interactions of syndecan-1 and syndecan-4 with alpha6beta4 integrin mediate human epidermal growth factor receptor (HER1 and HER2)-dependent motility and survival. J Biol Chem. 2014;289(44):30318–32.

17. McFall AJ, Rapraeger AC. Identification of an adhesion site within the syndecan-4 extracellular protein domain. J Biol Chem. 1997;272(20):12901–4.

18. McFall AJ, Rapraeger AC. Characterization of the high affinity cell-binding domain in the cell surface proteoglycan syndecan-4. J Biol Chem. 1998;273(43):28270–6.

19. Wang H, Jin H, Rapraeger AC. Syndecan-1 and Syndecan-4 Capture Epidermal Growth Factor Receptor Family Members and the alpha3beta1 Integrin Via Binding Sites in Their Ectodomains: NOVEL SYNSTATINS PREVENT KINASE CAPTURE AND INHIBIT alpha6beta4-INTEGRIN-DEPENDENT EPITHELIAL CELL MOTILITY. J Biol Chem. 2015;290(43):26103–13.

20. Serrano C, Cananzi S, Shen T, Wang LL, Zhang CL. Simple and highly specific targeting of resident microglia with adeno-associated virus. iScience. 2024;27(9):110706.

21. Hanlon KS, Cheng M, Ferrer RM, Ryu JR, Lee B, De La Cruz D, Patel N, Espinoza P, Santoscoy MC, Gong Y, et al. In vivo selection in non-human primates identifies AAV capsids for on-target CSF delivery to spinal cord. Mol Ther. 2024.

22. D’Costa S, Blouin V, Broucque F, Penaud-Budloo M, Francois A, Perez IC, Le Bec C, Moullier P, Snyder RO, Ayuso E. Practical utilization of recombinant AAV vector reference standards: focus on vector genomes titration by free ITR qPCR. Mol Ther Methods Clin Dev. 2016;5:16019.

23. Aurnhammer C, Haase M, Muether N, Hausl M, Rauschhuber C, Huber I, Nitschko H, Busch U, Sing A, Ehrhardt A, et al. Universal real-time PCR for the detection and quantification of adeno-associated virus serotype 2-derived inverted terminal repeat sequences. Hum Gene Ther Methods. 2012;23(1):18–28.

24. Ivanchenko MV, Hanlon KS, Devine MK, Tenneson K, Emond F, Lafond JF, Kenna MA, Corey DP, Maguire CA. Preclinical testing of AAV9-PHP.B for transgene expression in the non-human primate cochlea. Hear Res. 2020;394:107930.

